# Adaptation and serial choice bias are unaltered in autism

**DOI:** 10.1101/2021.09.13.460060

**Authors:** Ella Bosch, Matthias Fritsche, Christian Utzerath, Jan K. Buitelaar, Floris P de Lange

## Abstract

Autism Spectrum Disorder (ASD) or autism is characterized by social and non-social symptoms, including sensory hyper- and hyposensitivities. A suggestion has been put forward that some of these symptoms could be explained by differences in how sensory information is integrated with its context, including a lower tendency to leverage the past in the processing of new perceptual input. At least two history-dependent effects of opposite directions have been described in the visual perception literature: a repulsive adaptation effect, where perception of a stimulus is biased away from an adaptor stimulus, and an attractive serial choice bias, where perceptual choices are biased towards the previous choice. In this study, we investigated whether autistic participants differed in either bias from typically developing controls (TD). Sixty-four adolescent participants (31 with ASD, 33 TD) were asked to categorize oriented line stimuli in two tasks which were designed so that we would induce either adaptation or serial choice bias. Although our tasks successfully induced both biases, in comparing the two groups, we found no differences in the magnitude of adaptation nor in the modulation of perceptual choices by the previous choice. In conclusion, we find no evidence of a decreased integration of the past in visual perception of autistic individuals.

## Introduction

In typical perception, noisy sensory information is integrated with the spatial and temporal context in order to create a stable percept. In the case of temporal context, our environment tends to be temporally correlated or change in predictable ways. Because of this, perceptual systems, such as the visual system, can leverage the past in the processing of new sensory input (Schwartz et al., 2007). However, a consequence of this is that perception is biased by the past. Specifically, temporal context can bias current visual processing in two directions: a repulsive bias, known as an adaptation bias, and an attractive bias, known as a serial choice bias. Adaptation is a long-known and widely found phenomenon in which perception of a stimulus feature is biased away from the previous input (Kohn, 2007; Thompson & Burr, 2009; Webster, 2004, 2012, 2015). In contrast, serial choice bias, also known as sequential choice bias or choice repetition, is a phenomenon where the decision about a stimulus is biased towards the previous decision (Abrahamyan et al., 2016; Akaishi et al., 2014; Bosch et al., 2020; Braun et al., 2018; Fischer & Whitney, 2014; Fritsche et al., 2017; Fründ et al., 2014; St. John-Saaltink et al., 2016; Urai et al., 2017, 2019). These opposite biases may arise at different points of visual processing, with adaptation occurring during early stages of perception and serial choice bias occurring at later stages, possibly during decision-making (Bosch et al., 2020; Fritsche et al., 2017). Moreover, recent research suggests that they may be a reflection of distinct ways in which the visual system aims to optimize processing by increasing sensory sensitivity to changes of the environment while stabilizing percepts over time (Fischer & Whitney, 2014; Fritsche et al., 2020).

A suggestion that has been put forward is that autistic individuals may underutilize context in perceptual processing. Autism Spectrum Disorder (ASD) or autism is a developmental disorder that is most known for its social and behavioral symptoms, which feature prominently in the DSM-V diagnostic criteria. The behavioral symptoms also include sensory atypicalities, i.e. hyper- or hyporeactivity to sensory input or unusual interests in sensory aspects of the environment. In recent decades, different hypotheses have been formulated that attempt to explain these sensory atypicalities by how autistic individuals differ from typically developing (TD) individuals in the way that perceptual input is processed (e.g. Happé & Frith, 2006; Mottron et al., 2006; Pellicano & Burr, 2012). For instance, the Weak Central Coherence account (WCC; Happé & Frith, 2006) of autism conceptualizes a processing style that favors local processing over global, integrative processing, which may be observed as a reduction of the influence of the past in perceptual processing. Alternatively, work by Lawson et al. (2017) has found that autistic individuals overestimate the volatility of the environment, which could lead them to underutilize the past when processing new perceptual input. Whether due to a processing style or an overestimation of the volatility of the environment, if autistic individuals indeed underutilize temporal context, then they would be expected to show decreased biases that stem from this integration of context. Depending on where in the visual processing stream this occurs, they may show reduced adaptation, reduced serial choice bias, or both.

Evidence on adaptation in autism is mixed. Some research has indeed found evidence for reduced adaptation effects in autism using a variety of social and non-social visual stimuli, including faces (Ewing, Leach, et al., 2013; Ewing, Pellicano, et al., 2013; Pellicano et al., 2007), social eye-gaze (Lawson et al., 2017), biological motion (Karaminis et al., 2020; van Boxtel et al., 2016), and number (Turi et al., 2015). However, other studies have found no differences in adaptation to color (Maule et al., 2018) and causation (Karaminis et al., 2015). These differences in findings could be attributed to the use of different stimuli and designs, as well as differences in study population. Notably, there is some evidence that a higher severity of autistic traits and social atypicalities may be associated with larger reductions in the magnitude of adaptation (Lawson et al., 2017; Pellicano et al., 2007), suggesting there may be variation in adaptation decrease across the autistic spectrum.

Few studies have investigated the serial choice bias in autism. One study has found increased, rather than reduced, attractive influence of prior choices in visual location discrimination and visual-vestibular heading discrimination in autism (Feigin et al., 2021). However, another study found that perceptual decisions are less strongly attracted towards the immediate past in autistic individuals (Lieder et al., 2019). Although these studies both investigate the influence of the past, they do so by probing different biases using vastly different designs and stimuli. Moreover, these studies do not separate the influence of past stimuli from past decisions, complicating the interpretation of this work as reflecting a serial choice bias.

In summary, previous research on adaptation in autism has shown mixed results and serial choice bias in autism has hardly been investigated. Additionally, as no studies have looked into both biases and studies widely differ from each other with regards to their sample, design, and stimuli, it is difficult to compare findings between studies. This leaves open questions on how the past influences perception and perceptual decision making in autism.

In this study, we investigate whether autistic individuals differ from typically developing peers in the influence or use of prior information in perception and perceptual decision making. To this end, we conducted two psychophysical tasks in a sample of adolescent with and without ASD diagnosis. Both psychophysical tasks used the same type of line orientation stimuli, but were optimized in their design to induce either an adaptation bias or a serial choice bias. To preview the results, we indeed successfully induced both biases, but we found no differences between groups in the magnitude of their adaptation effect or the influence of the previous on the current choice. These findings suggest that integration of temporal context in visual processing of simple features in autism may be typical.

## Methods

### Data availability

All data and code used for stimulus presentation and analysis will be made available from the Donders Institute for Brain, Cognition and Behavior repository.

### Participants

The sample consisted of 64 participants (31 with ASD, 33 TD). Almost half of the sample was female (13 ASD, 16 TD). We tried to match participants in the groups as well as possible based on gender, age, and IQ.

The majority of participants with ASD were recruited from referrals to Karakter Child and Adolescent Psychiatry University Centre, Nijmegen, The Netherlands. The remainder of the participants were recruited through local schools, doctors offices, and recreational organizations such as sports clubs. Finally, some participants had previously participated in local studies and had given permission to be approached for other studies, and were thus recruited through local researchers (e.g. Utzerath et al., 2018, 2019).

All participants and their parent(s) or guardian provided written, informed consent. No parental consent was required for participants who were legal adults. Participants understood that they could withdraw from the study at any time. We compensated participants with gift vouchers. Participants were between 12 and 18 years old, native Dutch speakers, had normal or corrected-to-normal vision, and an IQ above 85. Exclusion criteria were (comorbid) major psychiatric or neurological disorders, current or recent alcohol or drug addiction, use of antipsychotic medication, claustrophobia, and pregnancy. An exception to the comorbid disorders was participants with an additional ADHD diagnosis, as this is a very frequent comorbidity (e.g. Jang et al., 2013). However, importantly, we included only participants of whom ASD was their primary diagnosis and who did not require ADHD medication. All participants in the ASD group had a clinical diagnosis of Autism Spectrum Disorder according to the DSM-5 criteria (American Psychiatric Association, 2013) or Autistic Disorder of Asperger’s Disorder according to the DSM-IV criteria (American Psychiatric Association, 1994). Additionally, we conducted a structured developmental interview (Autism Diagnostic Interview-Revised, ADI-R; Lord et al., 1994) to verify that their symptomatology matched the diagnostic threshold for ASD. In two cases, ASD diagnosis could not be confirmed and these participants were replaced. Members of the TD group had no history of neurological or psychiatric disorders. To screen for the presence of undiagnosed psychopathology, we conducted screening questionnaires (see below). Three participants were excluded based on these screening questionnaires, as they scored within the clinical range on the DSM-oriented scales. One additional TD participant was excluded due to receiving a developmental disorder diagnosis after participation in the study. Two participants were excluded from the ASD group due to their total IQ being under the preregistered cut-off (TIQ ≤ 85). Finally, two participants in the ASD group and two participants in the TD group were excluded due to poor performance on one of the experimental tasks. All excluded participants were replaced.

Recruitment and experimental procedures followed a protocol registered at and approved by and the local ethics committee (CCMO protocol NL60040.091.16, accessible at www.toetsingonline.nl).

### General procedure

All participants underwent the same general procedure. Participation consisted of a single experimental session and a set of questionnaires that could be completed during the session or at home. After providing written informed consent, we first conducted a brief IQ-test (see below) with the participant. If the participant was in the ASD group, we simultaneously conducted the structured interview (ADI-R; Lord et al., 1994) with their caregiver in a different room. Participants were then familiarized with the experimental setting in which they performed two behavioral tasks. The order of these tasks was fixed across participants due to an increase in response difficulty from the first to the second task. For each task, participants first received instructions, then performed practice blocks, and finally performed experimental blocks. This procedure was completed for the first task before introducing the second task. Participants were provided with several breaks during the session.

The IQ-test consisted of four subtests of the Dutch translation of the Wechsler Intelligence Scale for Children or Adults (WISC-III or WAIS-III; Kort et al., 2002; Wechsler, 1991; Wechsler et al., 2002) based on their age at inclusion. The subtests included were picture completion, similarities, block design, and vocabulary, in this order. In case a participant had already completed the WISC or WAIS (3^rd^ edition or later) within the two years before the inclusion date, for example as part of a clinical procedure or participation in a different scientific study, we did not conduct it again, as this would introduce retest effects, but instead requested and used their recent result.

The questionnaire set included, for all participants, Dutch translations of the self-report Edinburgh Handedness Inventory (Oldfield, 1971) and the Adolescent-Adult Sensory Profile (AASP; Brown & Dunn, 2002). Parents of TD participants completed the Child Behavior Checklist (CBCL; Achenbach, 1991) to control for the presence of psychopathology.

### Apparatus & stimuli

Visual stimuli were generated with the Psychophysics Toolbox (Brainard, 1997; Kleiner et al., 2007; Pelli, 1997) for MATLAB (2018) and displayed on a 24″ flat panel display (Benq XL2420T, resolution 1920 × 1080, refresh rate: 60 Hz). Participants viewed the stimuli from a distance of approximately 53 cm in a dimly lit room. A chinrest was used to ensure a constant viewing distance.

Orientation stimuli were generated by filtering white noise in the Fourier domain with a band-pass filter. The passband of spatial frequencies was defined as a Gaussian with a mean of 0.75 cycles/° and standard deviation of 0.3 cycles/°. The passband for orientations was defined as a von Mises distribution with location parameter µ and concentration parameter κ. The location parameter µ determined the mean orientation of a stimulus, while the concentration parameter κ effectively determined the amount of orientation noise. To introduce sensory uncertainty about the mean orientation of the stimulus we chose a low concentration parameter κ of 2.3, leading to uncertain stimuli containing multiple orientations around their mean orientation (see Figure 1a). After applying the inverse Fourier transform, the root mean square contrast of the stimuli was set to 11.76% of their mean luminance. All stimuli were windowed by a Gaussian envelope (2.3°s.d.). Stimuli as well as a white fixation dot were presented at the center of a grey background screen.

**Figure 1:**
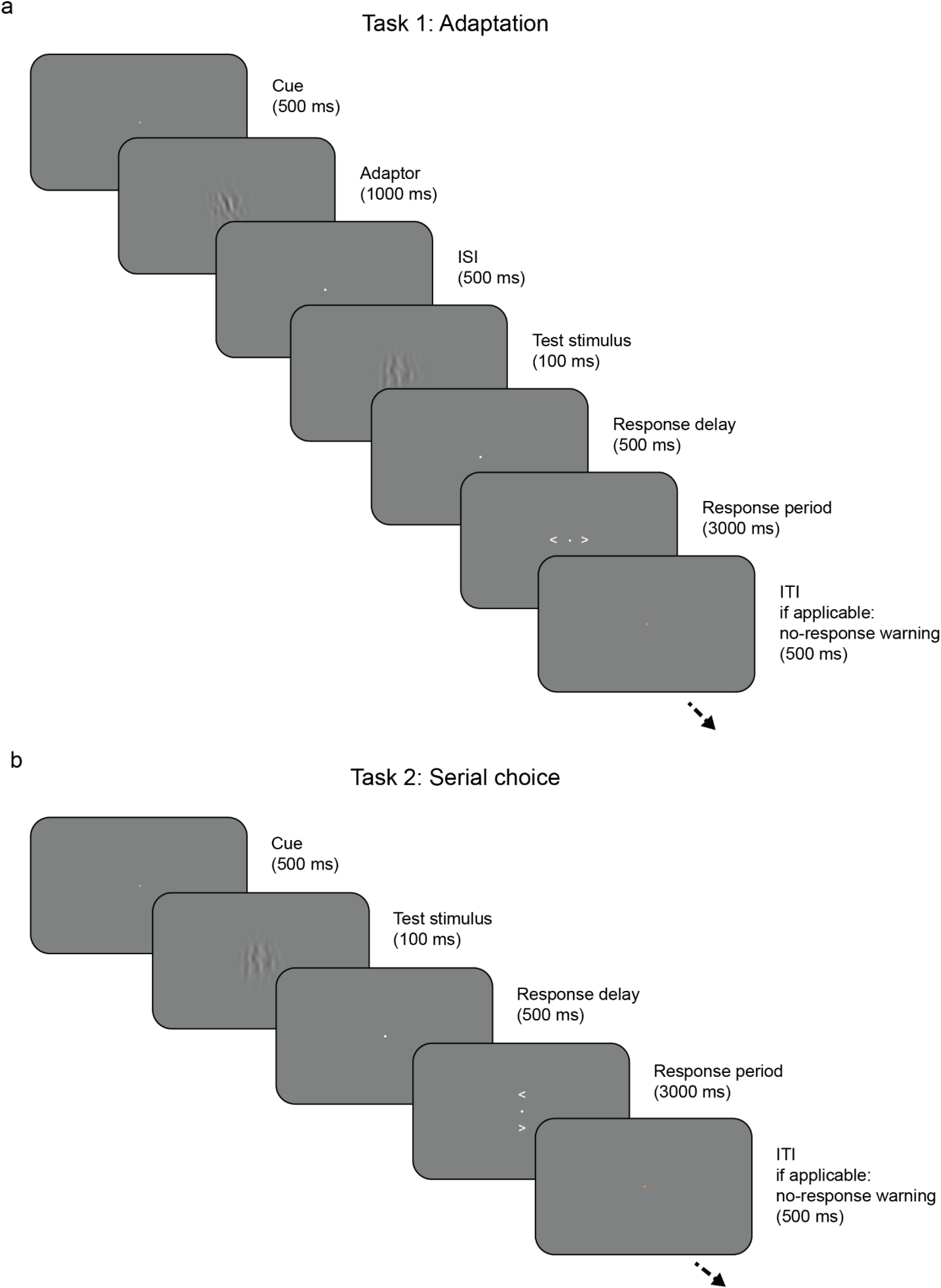
A single trial of task 1 (a) and task 2 (b). In both tasks, participants are presented ambiguous orientation stimuli and categorize the orientation of a test stimulus compared to vertical. Task 1 (a) is an adaptation task in which the test stimulus is preceded by an adaptor stimulus which participants are instructed is irrelevant. In task 2 (b), the serial choice task, the button mapping is pseudorandomized and communicated with a post-cue, after stimulus presentation.

In order to increase the participant’s interest in and engagement with the stimuli, we used a cover story in which the participant went on safari and was searching by the river for drinking zebras, represented by orientation stimuli, using an old spyglass, explaining the poor resolution of the visual stimuli. The stripes of the zebra were rotated away from vertical in either direction depending on which way the zebra was leaning to drink from the river.

### Task 1: Adaptation task

In each trial of task 1 (Figure 1a), two successive stimuli were presented on top of the fixation dot and separated by a 500 ms ISI. The first stimulus was oriented 20 degrees clockwise or counterclockwise from vertical, with each orientation equally frequent, and was presented for 1000 ms. We term this first stimulus the “adaptor”, as it was meant to induce a repulsive adaptation bias. The adaptor was instructed to be irrelevant (reeds growing by the river, according to the cover story) and had to be merely viewed. The second stimulus was oriented at or around vertical (−12, −6, 0, 6, or 12 degrees), with each orientation equally frequent, and was presented for 100 ms. We term this second stimulus the “test” stimulus, as it was meant to measure the biasing effect of the preceding adaptor. Participants were instructed to report the orientation of the test stimulus compared to vertical during the response period that followed the presentation of the test stimulus. During this response period, arrows pointing left and right were presented to the left and right of the fixation dot, respectively, for 3 seconds or until a response was given. Participants used the left and right arrow keys on the keyboard to give their answer, where the left and right keys indicated the stimulus was rotated counterclockwise (or “left” starting from the top) or counterclockwise (or “right” starting from the top), respectively. If participants did not respond within 3 seconds, the fixation dot would briefly turn orange to remind them to respond within the designated time.

Each practice block and experimental block consisted of 40 trials. Within each block, each combination of adaptor orientation and test stimulus orientation was equally likely and the order of trials was randomized. Participants completed at least one practice block and exactly four experimental blocks. After each block, participants received on-screen feedback on their performance on the easiest trials (i.e. the trials with the largest rotation away from vertical), though it was framed as general performance. Participants completed practice blocks until their performance on these easiest trials reached 75% (which was achieved after only one practice block for the majority of participants), with a reasonable distribution of left and right responses.

### Task 2: Serial choice task

Task 2 (Figure 1b) was similar to task 1, with the exception of the response configuration and the absence of an adaptor stimulus. In each trial of task 2, a single test stimulus was presented on top of the fixation dot. This stimulus was oriented at or around vertical (−5, 0, or 5 degrees), with the vertical orientation being the most frequent (occurring on 50% of trials). Participants were instructed to report the orientation of the test stimulus compared to vertical during the subsequent response period. During this response period, arrows pointing left and right were presented above and below the fixation dot for 3 seconds or until a response was given. Participants gave their answer using the up and down arrow keys on the keyboard, with the on-screen arrows indicating which key signified the direction of rotation (with left being rotated counter-clockwise or “left” starting from the top and right being rotated clockwise or “right” starting from the top). For this task, button mapping was pseudorandomized, with a 50% chance that the button mapping would flip between one trial and the next. If participants did not respond within 3 seconds, the fixation dot would briefly turn orange to remind them to respond within the designated time.

Each practice block consisted of 41 trials and each experimental block consisted of 81 trials. Within each block, each combination of test stimulus orientation and button mapping within a trial was equally likely. The order of trials was randomized in practice blocks and pseudorandomized within experimental blocks so that the frequency of stimulus orientation within successive trials (*t* and *t + 1*) was balanced as would be expected based on the frequency of each orientation. Participants completed at least one practice block and exactly three experimental blocks. After each block, participants received on-screen feedback on their performance, with vertical trials always being counted as correct as there was no correct answer on these trials. Participants completed practice blocks until their performance on these trials exceeded 75% (which was achieved after only one practice block for the majority of participants), with a reasonable distribution of left and right responses.

### Data cleaning

For both tasks, trials in which no response was given were removed from the data before analysis. In addition, for task 2, the serial choice task, trials with premature responses (≤ 200 ms from the onset of the button mapping) were removed. As a result, for task 1, the adaptation task, 36 out of 10,202 total trials (0.35%) were removed; for task 2, 115 of 15,552 total trials were removed (0.74%), of which 73 due to no response and 42 due to being below the response time cut-off. Trials removals from participants in the ASD group account for the majority of the removals: 63.9% and 70.4% of removed trials for task 1 and 2 respectively.

### ANALYSIS

For both tasks, response accuracy was calculated:

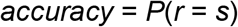

where *r* was the direction of the response and *s* the direction of the stimulus, with *s* constrained to be non-zero (i.e. non-vertical), leaving a counterclockwise vs clockwise binary for both variables.

We analyzed adaptation and choice repetition biases in three different ways: (1) by conditioning the current response on the preceding adaptor orientation or previous response (model-free analysis); (2) by fitting psychometric functions to the response data and quantifying shifts in the psychometric functions depending on preceding adaptor or previous response (psychometric analysis); (3) by fitting a hierarchical multiple logistic regression model to the data, accounting for the influence of both previous stimuli and responses (history-dependent multiple regression model).

### Model-free analysis

For task 1, we calculated the bias induced by the adaptor by calculating the difference in the proportion for clockwise responses for trials that had a counterclockwise adaptor versus trials that had a clockwise adaptor:

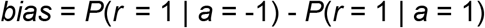

where *r* was the direction of the response and *a* the direction of the adaptor, with both variables representing either a counterclockwise (−1) or a clockwise (+1) direction.

For task 2, we then calculated the choice repetition probability as the mean of the probability to repeat a counterclockwise response and the probability to repeat a clockwise response:

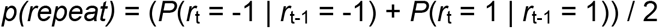

where *r*_*t*_ and *r*_*t-1*_ indicate the direction of the response on the current and previous trial. The probability was calculated per response direction to prevent a general response bias in either direction – in other words, a tendency to respond either counterclockwise or clockwise more often throughout the task – to influence *p(repeat)*.

Next, we conducted independent samples t-tests and Bayesian independent samples t-tests in JASP (JASP Team, 2020) to test for differences between the ASD group and the TD group in the calculated measures and quantify evidence in favor of and against the null hypotheses of no effect (no difference from 0 for bias, no difference from 0.5 for *p(repeat)*) or no difference between groups. We used default Cauchy priors (scale 0.707) for all Bayesian t-tests.

### Psychometric analysis

Next, we conducted an analysis that involved fitting a psychometric function to the data. Specifically, we employed a psychometric function fitting approach in order to quantify the effect of the adaptor on the response direction. As the limited number of trials per stimulus orientation did not allow for a good psychometric fit for all participants, we applied this method on pooled data of all participants within each group instead of on single-subject data.

We first expressed the probability of a clockwise response (*P*(*r*_*t*_ = 1)) as a function of the stimulus evidence 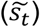 and fit a psychometric function (Figure 2a; Wichmann & Hill, 2001) of the form

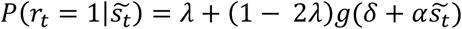

where *λ* was the probability of stimulus-independent errors (‘lapses’), *g* was the cumulative normal function, parameter *α* reflects perceptual sensitivity, and *δ* was a bias term corresponding to a general bias. The free parameters *λ, α* and *δ* were estimated by using the Palamedes toolbox for analyzing psychophysical data (Prins & Kingdom, 2018) using a maximum likelihood criterion.

**Figure 2:**
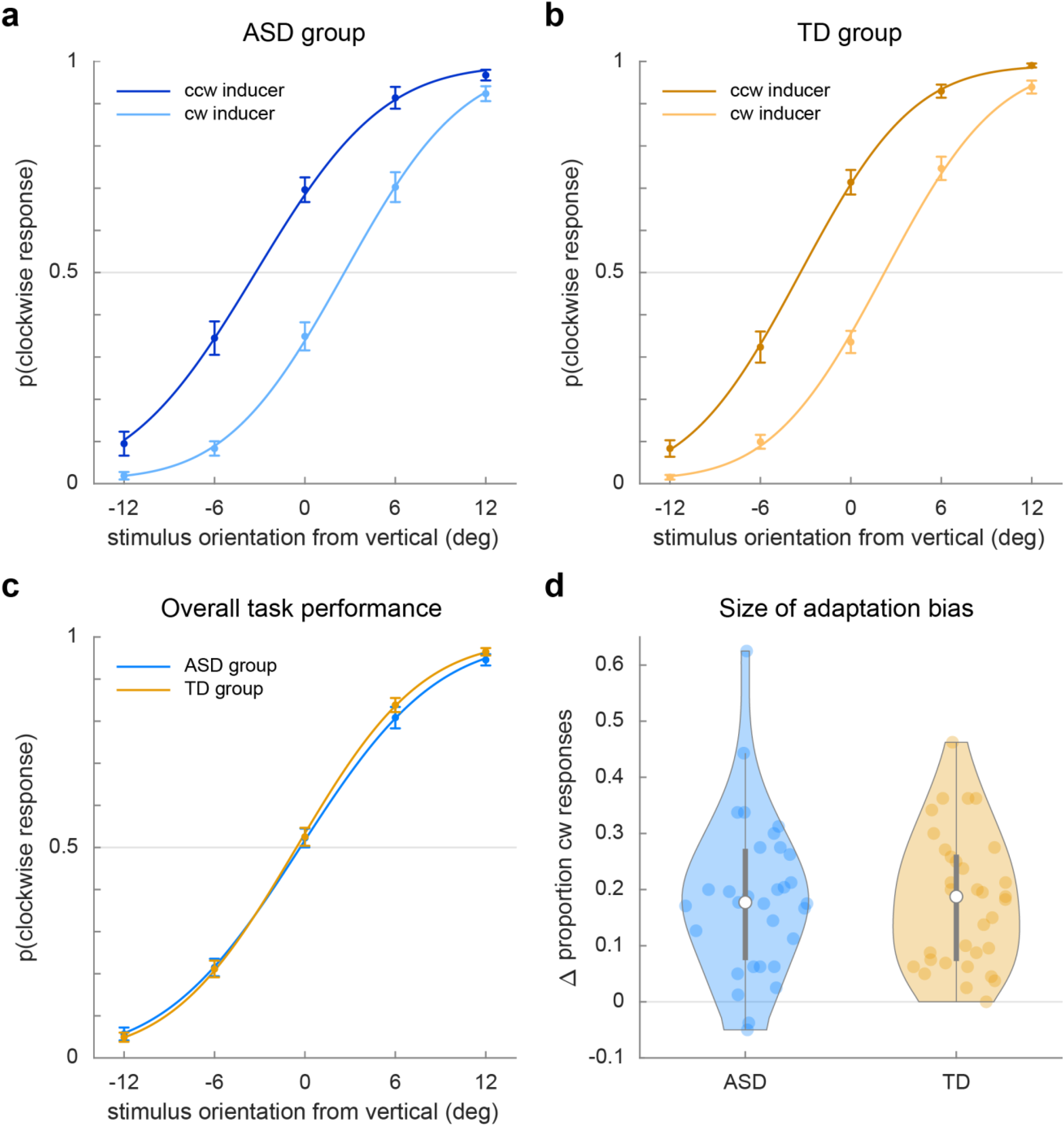
Adaptation task responses. (a) and (b) show the proportion of clockwise responses to stimuli following a counterclockwise (ccw) or clockwise (cw) adaptor for the ASD and TD group respectively. (c) shows the proportion of clockwise responses to stimuli, regardless of the direction of the adaptor, of the ASD group (blue) and the TD group (orange). (d) shows the magnitude of the adaptation bias, expressed as the difference in the proportion of clockwise responses after a clockwise or counterclockwise adaptor. Positive values indicate a repulsion away from the adaptor stimulus.

For the quantification of adaptation bias, we split the data into two bins based on the direction of the adaptor and then fit a psychometric curve to data in each bin. We then calculated the difference between the points of subjective equality (PSE) of each curve. This difference corresponds to the bias induced by the adaptor.

To test for differences between groups in lapse rate, perceptual sensitivity, general response bias, perceptual sensitivity, and bias induced by the adaptor, we used permutation tests. For each permutation, we randomly shuffled the group labels across participants thereby permuting ASD and TD assignments. We then applied the same psychometric fitting method described above. We repeated this method for 10,000 permutations. For each permutation, we computed the differences between groups of the lapse, perceptual sensitivity, and bias terms, as well as the bias induced by the adaptor. As p-values we report the percentage of permutations that led to more extreme values than those estimated on the empirical data. As we conducted a two-sided test, we multiplied this p-value by 2 and set the significance level to α = 0.05.

### History-dependent multiple regression model

The approaches described above allowed us to estimate biases induced by the adaptor (task 1) and previous decision (task 2) by splitting the data according to these variables. However, previous research (e.g. Bosch et al., 2020) has shown that this method of splitting data can partition meaningful variance and introduce or mask influences of other variables. For example, splitting by previous response can obscure a potential effect of the previous stimulus, which contributes to the serial choice patterns in the data of task 2. In order to estimate separate influences of different current-trial variables and previous-trial variables on the current decision, we constructed a generalized linear mixed model (GLMM) for task 2.

The GLMM contained a binomial link function to predict the current decision (counterclockwise or clockwise) based on the previous decision and other trial characteristics, as well as interactions between these factors. The factors in this regression model can be conceptually split into current-trial factors, history factors, and the group factor. The current-trial factors consist of the stimulus information (i.e. evidence direction) and button mapping on the current trial. The history factors consist of the stimulus information (i.e. evidence direction) and response characteristics (i.e. decision, button pressed, response time) of the previous trial. The group factor identifies the observer’s group. An overview of the GLMM with group factor can be found in Table 2 and is described below.

As we were interested in serial choice effects, we were interested in the influence of the previous decision on the current decision. Accordingly, we added the effect of previous decision (*pDecision*; clockwise or counterclockwise) as a factor to the model. In order to compare the effect of the previous decision with that of the stimulus information on the previous trial, we also added the identity (clockwise or counterclockwise from vertical) of the previous stimulus (*pStimIdent*). Next, to examine whether the influence of the previous decision or previous stimulus identity was modulated by the previous response time, we added interaction factors (*pDecision x pRt* and *pStimIdent x pRt*). Crucially, in order to investigate any group-difference between these effects, we added further interaction effects between all aforementioned factors and the *group* factor.

All factors described thus far reflect history effects. However, observers’ decisions were primarily based on the stimulus information present in the current trial. Therefore, we included the orientation of the stimulus on the current trial to the model (*cStimIdent*) and allowed for the influence of the current stimulus to be modulated by group (*cStimIdent x group*). To account for the possibility of a difference in general response bias between groups, we also added *group* as a single factor to the model (the group-independent general reponse bias was reflected by the intercept of the model).

Additionally, we added factors to account for effects of button- and/or motor preferences. First, to account for a preference for responding with one button over the other, and consequently for an effect of the button mapping on the perceptual decision, we added the button mapping as a factor to the model (*cButtonMapping*). Second, we added a factor to account for a possible motor repetition or alternation effect (*pButtonXcButtonMapping*). As with all other factors, we accounted for possible group differences in these effects by adding their interactions with *group* (*cButtonMapping x group* and *pButtonXcButtonMapping x group*).

Finally, we included the main effect of the previous response time (*pRt*) as well as its interaction with *group* (*pRt x group*). As these variables on their own provide no directional information, whether it be about the previous response or the stimulus information on the previous or current trial, they were unlikely to predict the decision on the current trial and were thus not expected to be significant factors in the model. We nevertheless added them to prevent that an unexpected modulation by these variables would show up in any of the interaction effects and hence be misinterpreted as such.

To investigate how variability in the strength of sensory atypicalities may affect perceptual decision-making, we constructed a second GLMM within the ASD group (N = 30; one subject was excluded due to missing AASP score). The factors included in this second GLMM were similar to those described above. The main difference was that the categorical *group* factor was replaced by a continuous *AASP* factor, which reflected the subjects’ AASP sum scores, both as a main factor and in all interactions that included *group*. See Table 3 for a full overview of the GLMM with AASP.

Before constructing the models, variables were (re-)coded as follows. Categorical predictors (*pDecision, pStimIdent, cStimIdent, pButton, cButtonMapping, pButtonXcButtonMapping*, and *group*) were coded using effect coding (−1/1). For *pDecision, pStimIdent, and cStimIdent*, −1 coded for the ccw direction and 1 for the cw direction. For *pButton*, −1 coded for the down button and 1 coded for the up button. For *cButtonMapping*, −1 coded for a configuration where the up button indicated the ccw direction and the bottom button indicated the cw direction, whereas 1 coded for the reverse configuration. For *pButtonXcButtonMapping*, a value of 1 indicated that pressing the same button as on the previous trial resulted in a cw response on the current trial, whereas −1 indicated that a repeated button press resulted in a ccw response. Finally, for *group*, the ASD group was coded as 1 and the TD group as −1. Non-categorical predictors were (re-)coded in the following ways. Response times (*pRt*) were transformed to robust z-scores by removing the subject-wise median and scaling the result by the subject-wise median absolute deviation (constant = 1.48). AASP scores were z-scored.

We used the R-package lme4 (Bates et al., 2015) to fit a generalized linear model from the binomial family. We fitted both models with ‘subjects’ as the only random grouping factor. For each fixed effect, we included its corresponding random slope coefficient, but without random correlations, as the model did not converge.

For significance testing we report Walds-Z test. Walds Z-test is valid only in the asymptotic regime assuming a multivariate normal sampling distribution of parameters and a proportional sampling distribution of the log likelihood to *χ*^2^. Therefore, we must be very conservative in our interpretation of the reported *p*-values if the effects are not obvious from effect-sizes alone. An overview of the model outputs can be found in Table 2 and Table 3.

## Results

### Sample characteristics

An overview of sample characteristics can be found in Table 1. The ASD group and TD group were comparable with regards to gender (*χ*^*2*^ = 0.277, *p* = 0.599). On average, participants were aged 15 years and 7 months on the day of inclusion, with no age difference between groups (*t*(62) = .209, *p* = .835). On average, the TD group scored higher on the Wechsler Intelligence Scale than the ASD group, which is a common occurrence in ASD literature. Specifically, there was a 7.94 point difference on the total scale (TIQ: *t*(62) = 2.794, *p* = 0.007), 9.47 point difference on the performance scale (PIQ: *t*(62) = 3.536, *p* = 0.001), and a 5.89 point difference on the verbal scale, although the last difference did not reach significance (VIQ: *t*(62) = 1.784, *p* = 0.079).

**Table 1:**
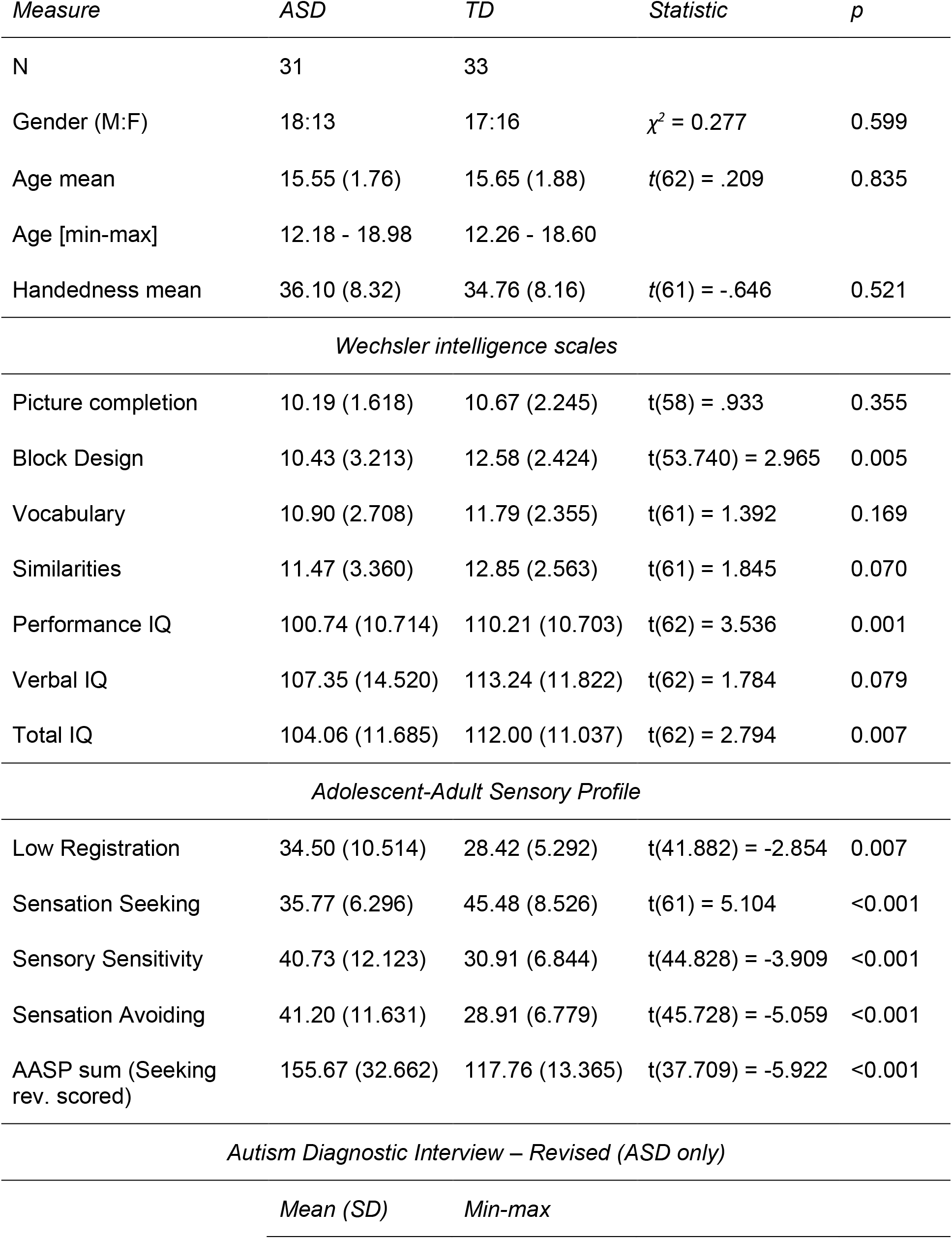
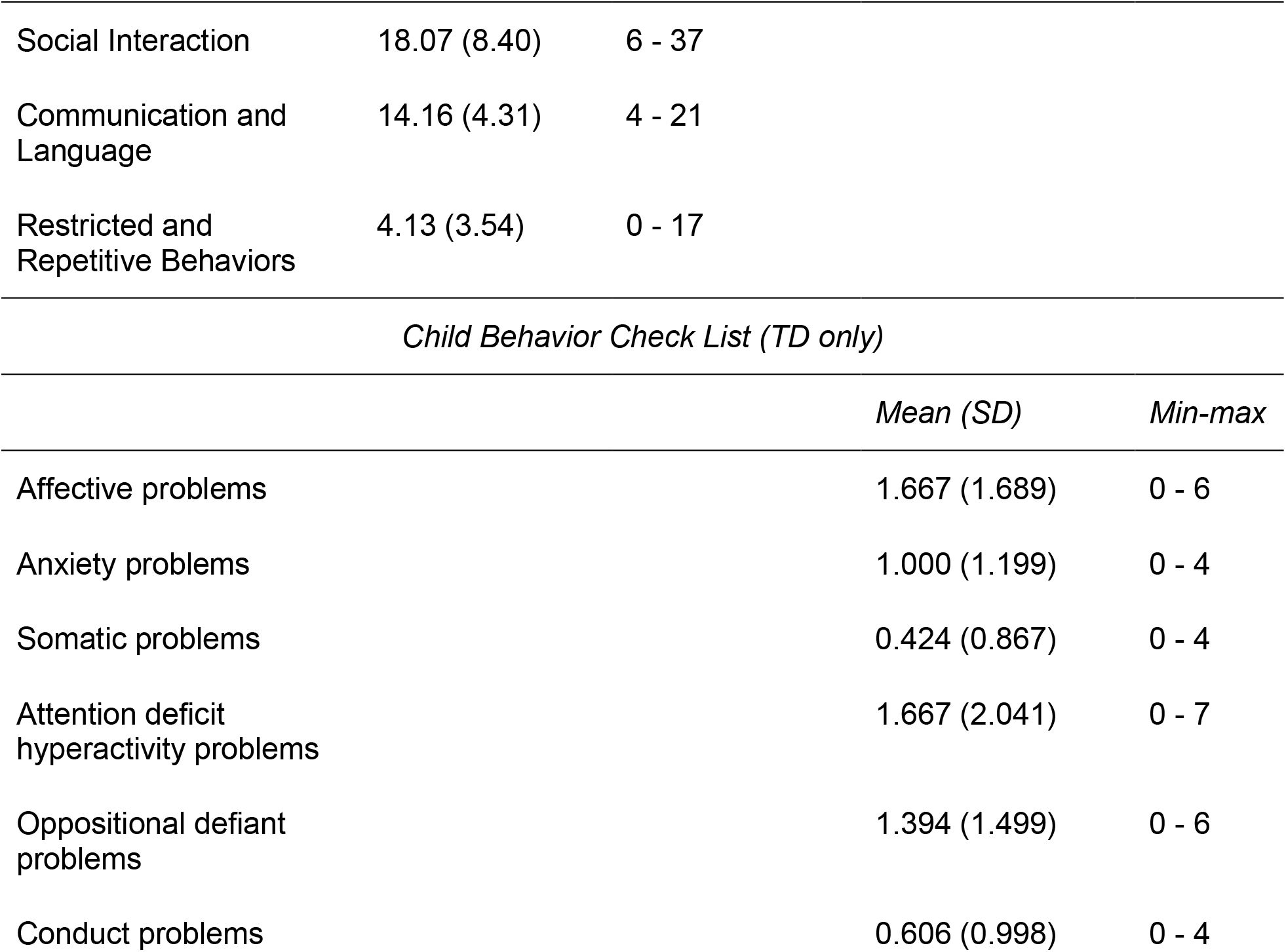
Sample characteristics.

| *Measure* | *ASD* | *TD* | *Statistic* | *p* |
| --- | --- | --- | --- | --- |
| N | 31 | 33 |  |  |
| Gender (M:F) | 18:13 | 17:16 | $\chi^2 = 0.277$ | 0.599 |
| Age mean | 15.55 (1.76) | 15.65 (1.88) | $t(62) = .209$ | 0.835 |
| Age [min-max] | 12.18 - 18.98 | 12.26 - 18.60 |  |  |
| Handedness mean | 36.10 (8.32) | 34.76 (8.16) | $t(61) = -.646$ | 0.521 |
| *Wechsler intelligence scales* |  |  |  |  |
| Picture completion | 10.19 (1.618) | 10.67 (2.245) | $t(58) = .933$ | 0.355 |
| Block Design | 10.43 (3.213) | 12.58 (2.424) | $t(53.740) = 2.965$ | 0.005 |
| Vocabulary | 10.90 (2.708) | 11.79 (2.355) | $t(61) = 1.392$ | 0.169 |
| Similarities | 11.47 (3.360) | 12.85 (2.563) | $t(61) = 1.845$ | 0.070 |
| Performance IQ | 100.74 (10.714) | 110.21 (10.703) | $t(62) = 3.536$ | 0.001 |
| Verbal IQ | 107.35 (14.520) | 113.24 (11.822) | $t(62) = 1.784$ | 0.079 |
| Total IQ | 104.06 (11.685) | 112.00 (11.037) | $t(62) = 2.794$ | 0.007 |
| *Adolescent-Adult Sensory Profile* |  |  |  |  |
| Low Registration | 34.50 (10.514) | 28.42 (5.292) | $t(41.882) = -2.854$ | 0.007 |
| Sensation Seeking | 35.77 (6.296) | 45.48 (8.526) | $t(61) = 5.104$ | <0.001 |
| Sensory Sensitivity | 40.73 (12.123) | 30.91 (6.844) | $t(44.828) = -3.909$ | <0.001 |
| Sensation Avoiding | 41.20 (11.631) | 28.91 (6.779) | $t(45.728) = -5.059$ | <0.001 |
| AASP sum (Seeking rev. scored) | 155.67 (32.662) | 117.76 (13.365) | $t(37.709) = -5.922$ | <0.001 |
| *Autism Diagnostic Interview – Revised (ASD only)* |  |  |  |  |
|  | *Mean (SD)* | *Min-max* |  |  |

|  |  |  |
| --- | --- | --- |
| Social Interaction | 18.07 (8.40) | 6 - 37 |
| Communication and Language | 14.16 (4.31) | 4 - 21 |
| Restricted and Repetitive Behaviors | 4.13 (3.54) | 0 - 17 |

*Child Behavior Check List (TD only)*
|  | *Mean (SD)* | *Min-max* |
| --- | --- | --- |
| Affective problems | 1.667 (1.689) | 0 - 6 |
| Anxiety problems | 1.000 (1.199) | 0 - 4 |
| Somatic problems | 0.424 (0.867) | 0 - 4 |
| Attention deficit hyperactivity problems | 1.667 (2.041) | 0 - 7 |
| Oppositional defiant problems | 1.394 (1.499) | 0 - 6 |
| Conduct problems | 0.606 (0.998) | 0 - 4 |

The ASD group and TD group differed significantly with regards to their sensory symptomatology as measured by the AASP. The ASD group self-reported higher scores on the subscales Low Registration (*t*(41.882) = −2.854, *p* = 0.007), Sensory Sensitivity (*t*(44.828) = −3.909, *p* < 0.001), and Sensation Avoiding (*t*(45.728) = −5.059, *p* < 0.001), and lower scores on the subscale Sensation Seeking (*t*(61) = 5.104, *p* < 0.001). A sum score was calculated by adding the values for each subscale, with the subscale Sensation Seeking reversed scored. The groups differed significantly on this sum score as well (*t*(37.709) = −5.922, *p* < 0.001).

The ASD diagnoses for participants in the ASD group were confirmed by the ADI-R (see Supplemental table 1 for individual scores). TD participants scored within normal range on the CBCL.

### Adaptation task

First, we established that participants were able to discriminate counterclockwise from clockwise stimuli in task 1 (adaptation task). Both the ASD group and TD group were well able to discriminate between counterclockwise and clockwise stimuli in task 1 (Figure 2c; mean performance of 87.1 ± 7.9% [65.6 - 96.1% range] in the ASD group and 88.6 ± 5.2% [74.8 - 95.3% range] in the TD group). We found no differences in overall response accuracy on the adaptation task between the ASD and TD groups (*accuracy*: t(62) = 0.924, *p* = 0.359; BF_10_ = 0.367, error % = 3.898e-4). Average response times were fast (ASD: mean = 408.3 ms, SD = 156.5 ms; TD: mean = 434.1 ms, SD = 197.4 ms) and not different between groups (*t*(62) = −0.5761, *p* = 0.567; BF_10_ = 0.294, error % = 0.002).

Next, we looked at whether the direction of the adaptor influenced participants’ responses. We observed a clear effect of the adaptor in both groups (Figure 2a and b), with the probability of a clockwise response after a counterclockwise versus clockwise adaptor increasing numerically for all but two participants in the ASD group and all but one participant in the TD group. The induced bias was statistically significant in both groups (Figure 2d: ASD: *t*(30) = 7.407, *p* < 0.001; BF_10_ = 4.66e5, error % = 1.19e-8; TD: *t*(32) = 8.734, *p* < 0.001; BF_10_ = 1.98e7, error % = 5.74e-10). On average, the proportion of clockwise responses differed between adaptor conditions by 18.7 ± 14.1%p in the ASD group and 18.1 ± 11.9%p in the TD group.

Finally, we looked at whether the influence of the adaptor was altered in ASD by comparing the magnitude of the induced bias between the ASD and TD group. We found that the magnitude of the bias did not differ between groups, with moderate evidence for a lack of difference (Figure 2d: *t*(63) = −0.206, *p* = 0.837; BF_10_ = 0.260, error % = 0.002). This suggests that the repulsive adaptation bias was similar across groups.

Our psychometric fitting approach with permutation testing showed similar results. We found no difference between the ASD group and TD group in overall bias (*p* = 0.6152), slope (*p* = 0.6724), lapses (*p* = 0.6298), and bias induced by the adaptor (*p* = 0.7086), suggesting that the groups exhibited similarly large adaptation biases.

### Serial choice task

For the serial choice task, we first established that participants were able to discriminate counterclockwise from clockwise stimuli. Both the ASD group and TD group were able to do this discrimination, (Figure 3c: mean response accuracy of 77.0 ± 11.2% [55.9 - 95.0% range] in the ASD group and 80.7 ± 9.3% [60.7 - 93.3% range] in the TD group). We found no significant differences in response accuracy on this serial choice task between the groups (*accuracy*: t(62) = 1.432, *p* = 0.157; BF_10_ = 0.606, error % = 0.003). Mean response times did not differ between groups (ASD: mean = 1039.4 ms, SD = 195.3 ms; TD: mean = 1065.3 ms, SD = 157.9 ms; *t*(62) = −0.5831, *p* = 0.5619; BF_10_ = 0.295, error % = 0.001).

**Figure 3:**
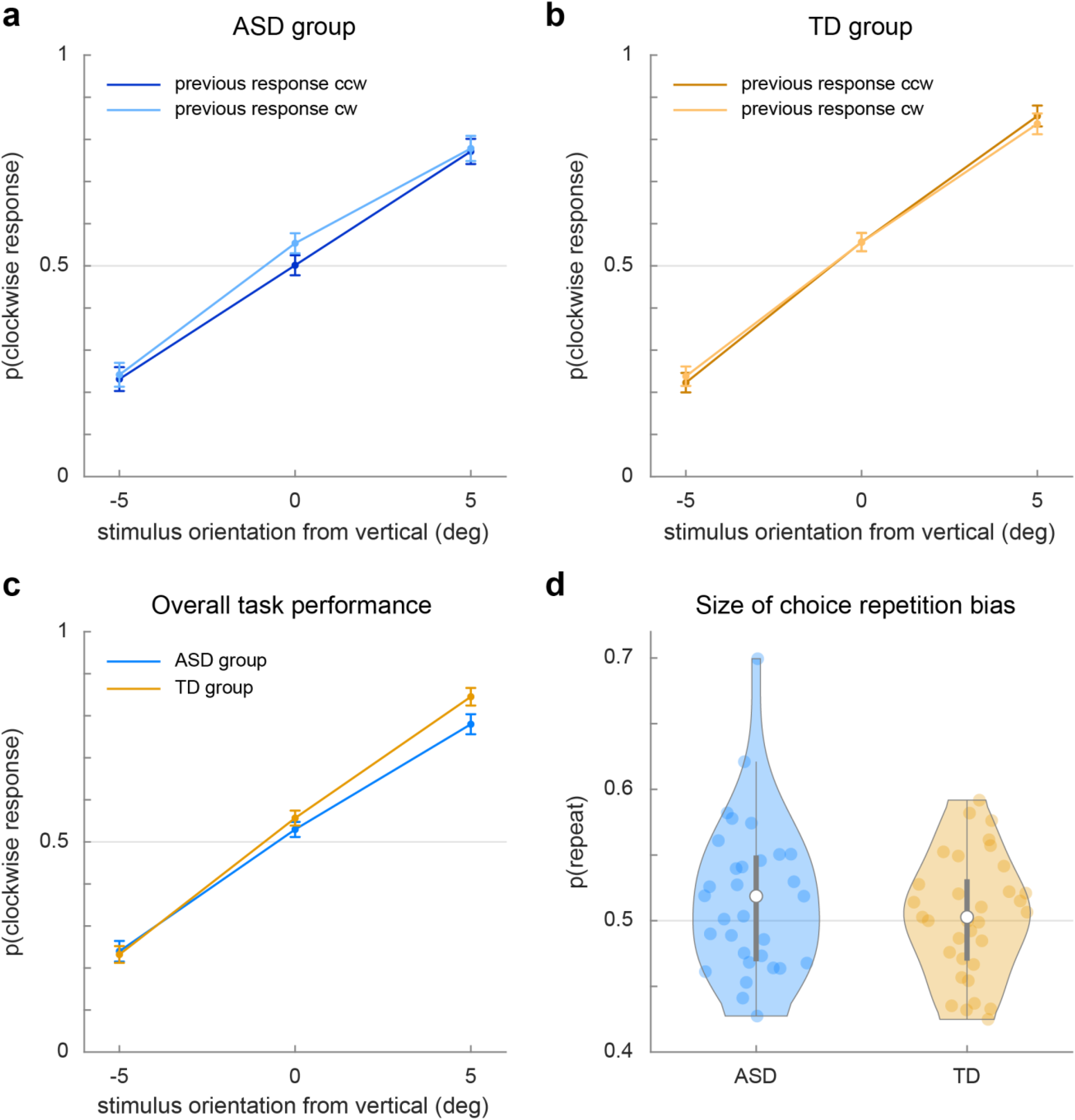
Serial choice task responses. (a) and (b) show the proportion of clockwise responses to stimuli after a counterclockwise (ccw) or clockwise (cw) response on the previous trial, for the ASD and TD group respectively. (c) shows the proportion of clockwise responses to stimuli, regardless of the previous response, of the ASD group (blue) and the TD group (orange). (d) shows the magnitude of the choice repetition probability (*p(repeat)*).

Next, we looked at whether participants’ previous choice influenced their current choice. We did not observe an apparent effect of the previous choice in either group (Figure 3a and b). We quantified the effect by calculating the probability that participants repeated the previous choice (choice repetition). On average, choice repetition probability was 51.7% ± 5.8% [42.7 – 69.9% range] in the ASD group and 50.3% ± 4.6% [42.5 – 59.2% range] in the TD group (Figure 3d). For neither group, this probability differed convincingly from chance (ASD: *t*(30) = 1.642, *p* = 0.111; BF_10_ = 0.637, error % = 0.008; TD: *t*(32) = 0.380, *p* = 0.707; BF_10_ = 0.199, error % = 1.90e-6). Although we hypothesized differences in choice repetition probability between the groups, we did not find this (*t*(62) = −1.077, *p* = 0.286), nor did we find convincing evidence against the null hypothesis (BF_10_ = 0.417, error % = 1.456e-4).

As we set out to investigate how the previous choice influences the current choice and how this may differ between autistic and non-autistic individuals, the fact that we did not find a serial choice bias may seem problematic. However, research in a typical population has shown that serial choice bias may be obscured across trials by simultaneous but oppositely signed effects, for example a repulsive effect of the stimulus of the previous trial (sensory adaptation) and an attractive effect of the response on said trial (Bosch et al., 2020). Therefore, in order to study the effect of the previous response while controlling for concurrent stimulus-related effects, we used an analytical method that can identify these separate effects. We applied a GLMM method for this reason.

Indeed, the GLMM (Figure 4; see Table 2 for a full model overview) revealed a small yet reliable attractive effect of the previous decision on the current decision (*pDecision*: *b* = 0.0686, *SE* = 0.0328, *p* = 0.036). We also found that people were more likely to repeat fast trials (*pDecision x pRT*: *b* = −0.0622, *SE* = 0.0156, *p* = 6.56e-5). Both effects have been found in a previous study using a comparable analysis (Bosch et al., 2020). Interestingly, the repulsive effect of the previous stimulus on the current decision was not replicated in this dataset (*pStimIdent*: *b* = −0.0102, *SE* = 0.0346, *p* = 0.769), perhaps because the orientation information in the current stimuli was much weaker and noisier, reducing or even removing the influence of this information on the current decision.

**Figure 4:**
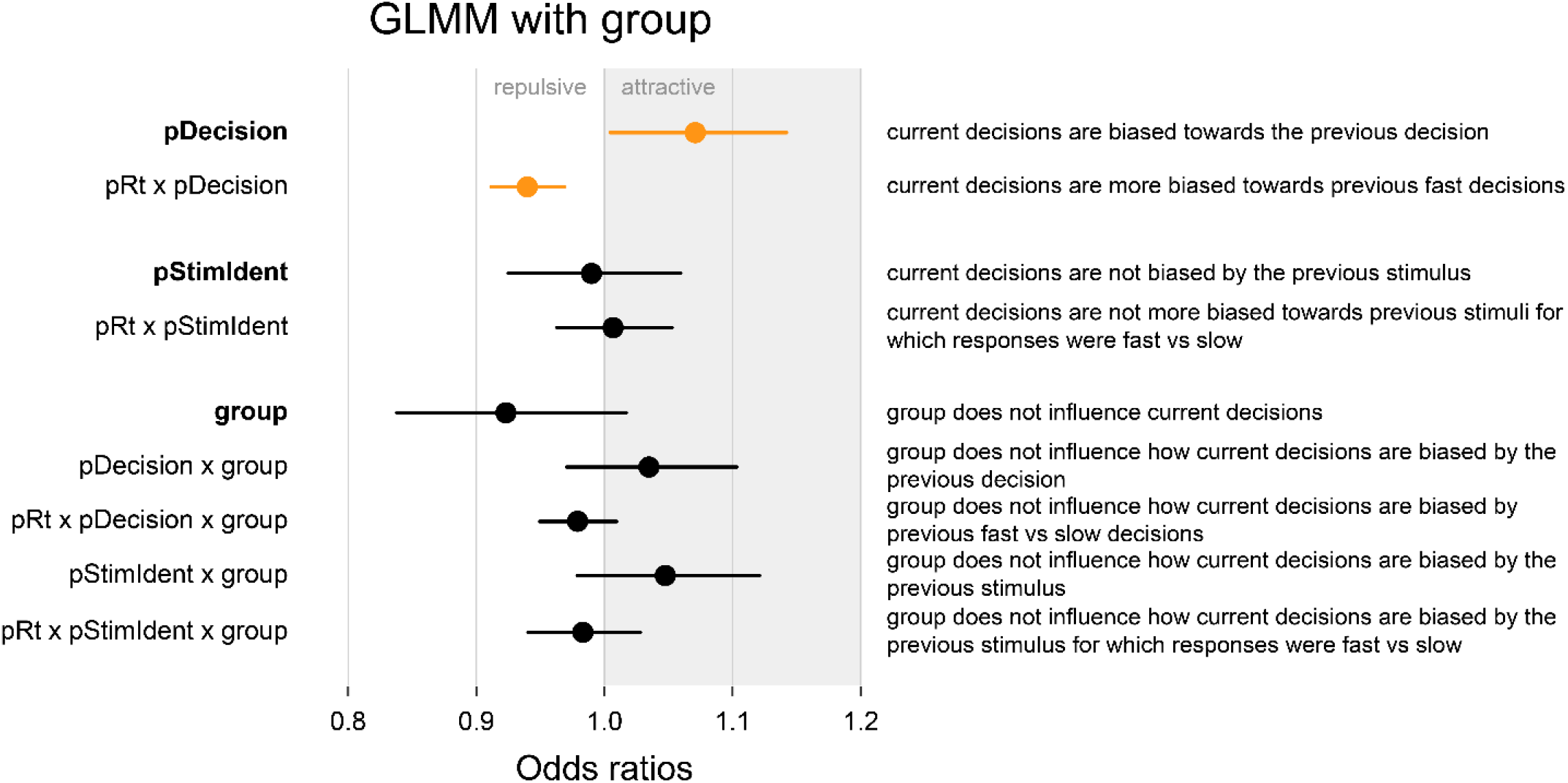
GLMM results for a model over all participants (N = 64) that predicts the current decision based on current- and previous trial factors and group (ASD vs TD). Not all factors are shown (see table 2 for full model overview); significant factors are marked in orange. Results show that decisions are biased towards the previous decision, and more biased towards previous fast decisions than slow decisions. No group differences were found.

**Table 2:**
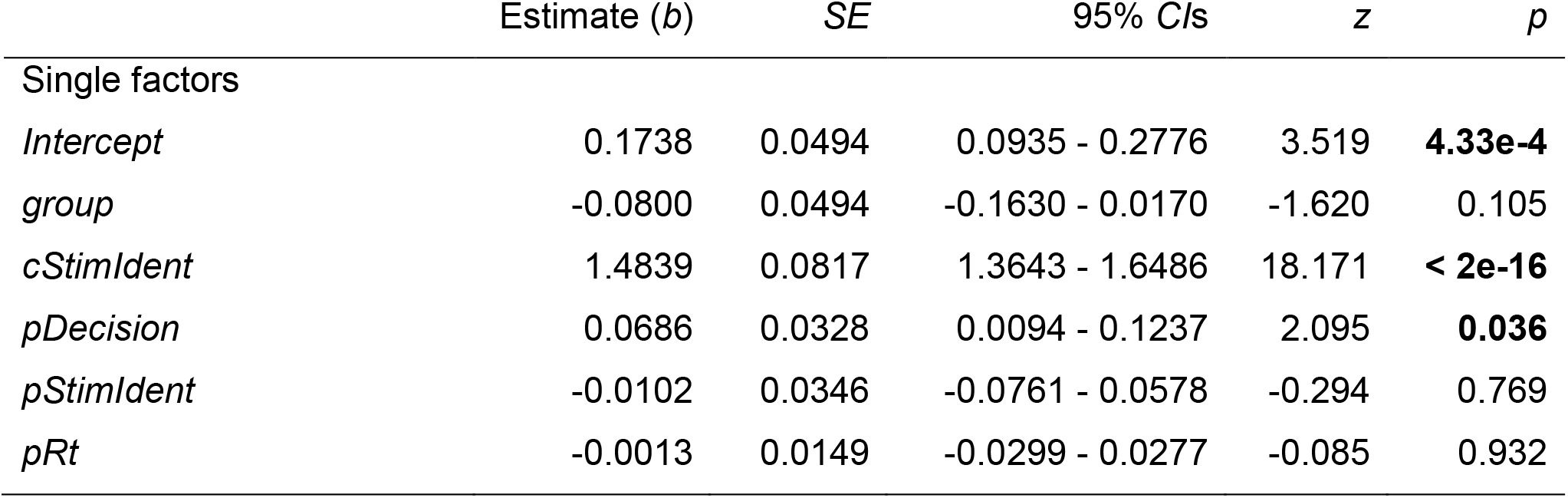
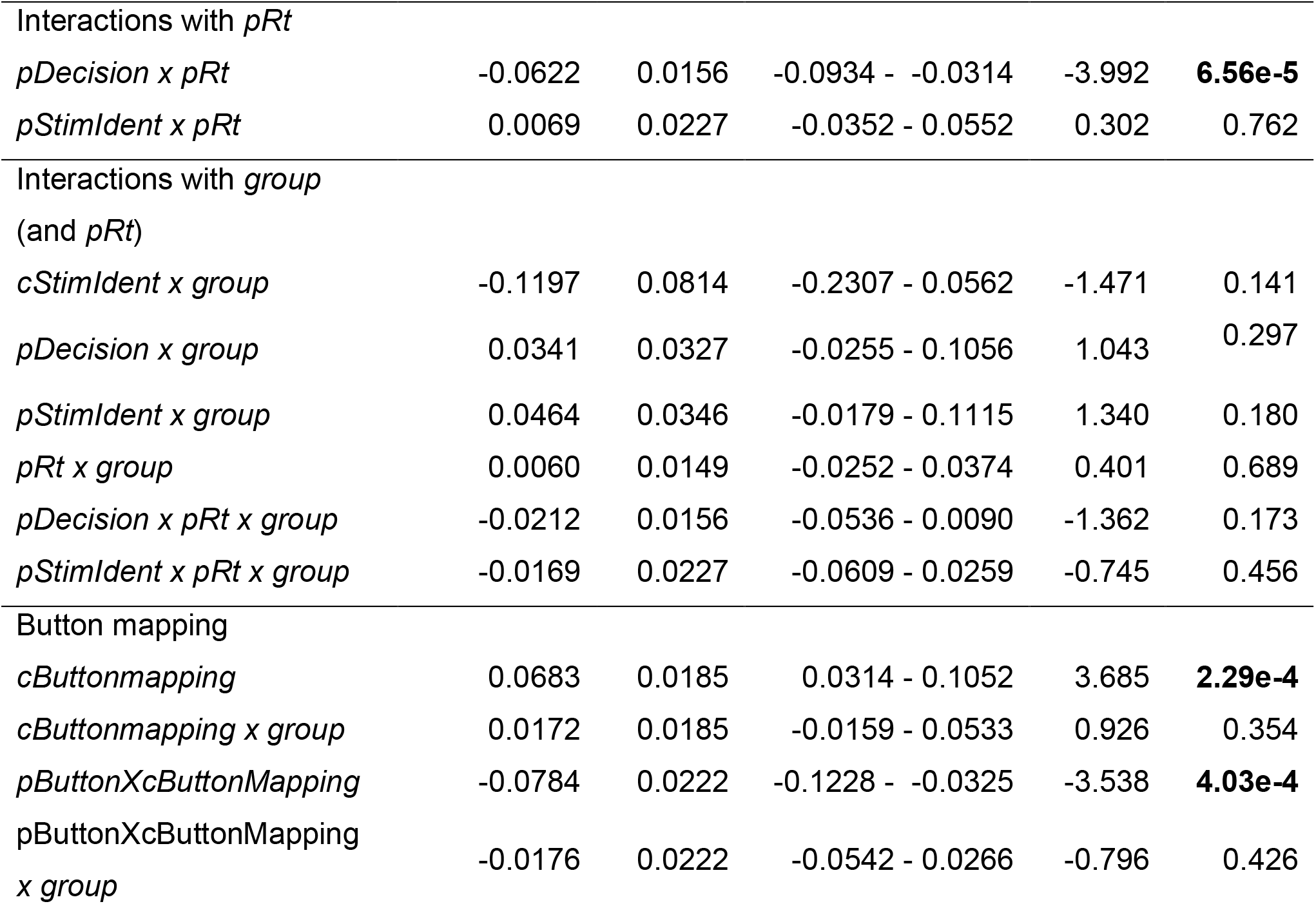
GLMM fixed factors for a model over all participants (N = 64) that predicts the current decision based on current- and previous trial factors and group (ASD vs TD).

|  | Estimate (*b*) | SE | 95% CIs | *z* | *p* |
| --- | --- | --- | --- | --- | --- |
| Single factors |  |  |  |  |  |
| *Intercept* | 0.1738 | 0.0494 | 0.0935 - 0.2776 | 3.519 | **4.33e-4** |
| *group* | -0.0800 | 0.0494 | -0.1630 - 0.0170 | -1.620 | 0.105 |
| *cStimIdent* | 1.4839 | 0.0817 | 1.3643 - 1.6486 | 18.171 | **< 2e-16** |
| *pDecision* | 0.0686 | 0.0328 | 0.0094 - 0.1237 | 2.095 | **0.036** |
| *pStimIdent* | -0.0102 | 0.0346 | -0.0761 - 0.0578 | -0.294 | 0.769 |
| *pRt* | -0.0013 | 0.0149 | -0.0299 - 0.0277 | -0.085 | 0.932 |
| Interactions with *pRt* |  |  |  |  |  |
| *pDecision x pRt* | -0.0622 | 0.0156 | -0.0934 - -0.0314 | -3.992 | **6.56e-5** |
| *pStimIdent x pRt* | 0.0069 | 0.0227 | -0.0352 - 0.0552 | 0.302 | 0.762 |
| Interactions with *group*<br>(and *pRt*) |  |  |  |  |  |
| *cStimIdent x group* | -0.1197 | 0.0814 | -0.2307 - 0.0562 | -1.471 | 0.141 |
| *pDecision x group* | 0.0341 | 0.0327 | -0.0255 - 0.1056 | 1.043 | 0.297 |
| *pStimIdent x group* | 0.0464 | 0.0346 | -0.0179 - 0.1115 | 1.340 | 0.180 |
| *pRt x group* | 0.0060 | 0.0149 | -0.0252 - 0.0374 | 0.401 | 0.689 |
| *pDecision x pRt x group* | -0.0212 | 0.0156 | -0.0536 - 0.0090 | -1.362 | 0.173 |
| *pStimIdent x pRt x group* | -0.0169 | 0.0227 | -0.0609 - 0.0259 | -0.745 | 0.456 |
| Button mapping |  |  |  |  |  |
| *cButtonmapping* | 0.0683 | 0.0185 | 0.0314 - 0.1052 | 3.685 | **2.29e-4** |
| *cButtonmapping x group* | 0.0172 | 0.0185 | -0.0159 - 0.0533 | 0.926 | 0.354 |
| *pButtonXcButtonMapping* | -0.0784 | 0.0222 | -0.1228 - -0.0325 | -3.538 | **4.03e-4** |
| *pButtonXcButtonMapping x group* | -0.0176 | 0.0222 | -0.0542 - 0.0266 | -0.796 | 0.426 |

We then looked at whether the effects involving the previous decision were modulated by group and found that they were not (*pDecision x group*: *b* = 0.0341, *SE* = 0.0327, *p* = 0.297; *pDecision x pRT x group*: *b* = −0.0212, *SE* = 0.0156, *p* = 0.173). This suggests that group does not alter the effect of the previous decision on the previous trial, nor does it alter the modulation of this effect by previous response time.

The failure to find group differences in our data may be due to heterogeneity within the ASD and typical population. ASD is described as a spectrum, with social, behavioral, but also sensory characteristics varying between individuals on this spectrum. It is possible that investigating the magnitude of choice repetition bias not between diagnostic groups but along the dimension of sensory atypicality may reveal an effect of this symptomatology specifically. For this reason, we created a separate GLMM within the ASD group and included the sum score on the AASP as a factor (Figure 5; see Table 3 for a full model overview). As in the previous model, observers were more likely to repeat a previous decision, although this effect did not reach significance (*pDecision*: *b* = 0.0970, *SE* = 0.0519, *p* = 0.062), and observers were more likely to repeat previous fast decisions (*pDecision x pRt*: *b* = −0.0829, *SE* = 0.0226, *p* = 2.44e-4). There was a lower tendency of observers with stronger sensory atypicalities (reflected by high AASP scores) to repeat previous decisions, although this effect did not reach significance (*pDecision x AASP*: *b* = −0.0862, *SE* = 0.0515, *p* = 0.094). Moreover, observers with stronger sensory atypicalities were less likely to repeat fast decisions (*pDecision x pRt x AASP*: *b* = 0.0449, *SE* = 0.0219, *p* = 0.040). However, we found the model predictions from this model did not closely fit the data except for participants with AASP scores closest to the mean, bringing into question the reliability of the model results. We therefore choose to remain cautious in our interpretation of these findings and emphasize that these potential subtle effects require replication in future studies.

**Figure 5:**
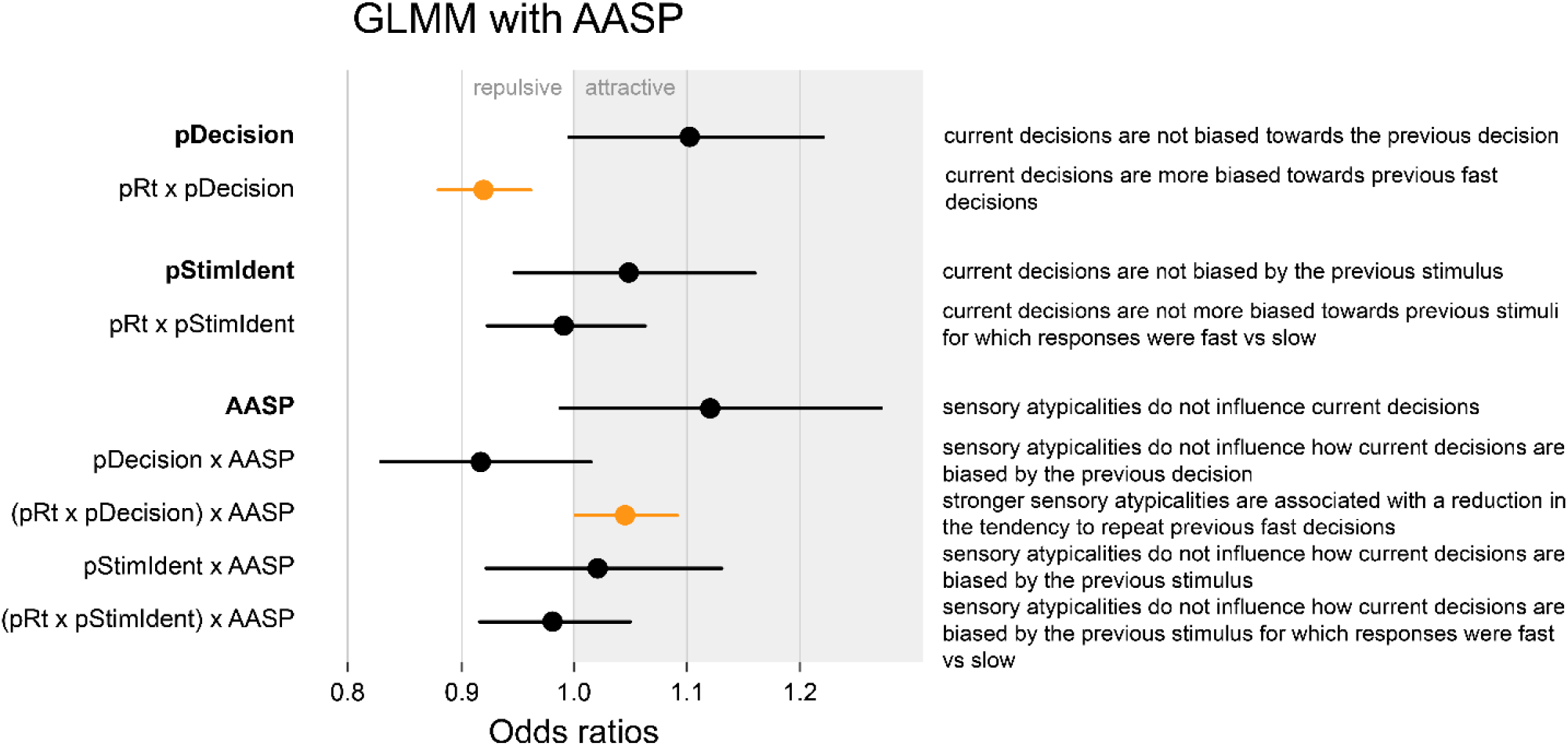
GLMM results for a model within the ASD group (N = 30) that predicts the current decision based on current- and previous trial factors and the participant’s AASP sum score. Not all factors are shown (see Table 3 for full model overview); significant factors are marked in orange. Results show that decisions are more biased towards fast than slow previous decisions, and that this bias is weaker for participants with higher AASP sum scores.

**Table 3:** GLMM fixed factors for a model within the ASD group (N = 30) that predicts the current decision based on current- and previous trial factors and the participant’s AASP sum score.

|  | Estimate (*b*) | SE | 95% CIs | *z* | *p* |
| --- | --- | --- | --- | --- | --- |
| Single factors |  |  |  |  |  |
| *Intercept* | 0.0967 | 0.0657 | -0.0125 - 0.2307 | 1.472 | 0.141 |
| *AASP* | 0.1171 | 0.0648 | 0.0163 - 0.2451 | 1.808 | 0.071 |
| *cStimIdent* | 1.3748 | 0.1274 | 1.1859 - 1.6140 | 10.791 | **< 2e-16** |
| *pDecision* | 0.0970 | 0.0519 | -0.0072 - 0.2103 | 1.869 | 0.062 |
| *pStimIdent* | 0.0473 | 0.0518 | -0.0355 - 0.1509 | 0.912 | 0.362 |
| *pRt* | 0.0063 | 0.0218 | -0.0319 - 0.0469 | 0.287 | 0.774 |
| Interactions with *pRt* |  |  |  |  |  |
| *pDecision x pRt* | -0.0829 | 0.0226 | -0.1261 - -0.0377 | -3.669 | **2.44e-4** |
| *pStimIdent x pRt* | -0.0093 | 0.0358 | -0.0747 - 0.0602 | -0.261 | 0.794 |
| Interactions with AASP |  |  |  |  |  |
| sum (and pRt) |  |  |  |  |  |
| *cStimIdent x AASP* | 0.1603 | 0.1255 | -0.0380 - 0.4131 | 1.278 | 0.201 |
| *pDecision x AASP* | -0.0862 | 0.0515 | -0.1811 - 0.0173 | -1.673 | 0.094 |
| *pStimIdent x AASP* | 0.0211 | 0.0521 | -0.0697 - 0.1206 | 0.406 | 0.685 |
| *pRt x AASP* | -0.0079 | 0.0211 | -0.0489 - 0.0330 | -0.374 | 0.708 |
| *pDecision x pRt x AASP* | 0.0449 | 0.0219 | -0.0068 - 0.0896 | 2.050 | **0.040** |
| *pStimIdent x pRt x AASP* | -0.0120 | 0.0345 | -0.0879 - 0.0475 | -0.577 | 0.564 |
| Button mapping |  |  |  |  |  |
| *cButtonmapping* | 0.0854 | 0.0283 | 0.0338 - 0.1393 | 3.019 | **2.53e-3** |
| *pButtonXcButtonMapping* | -0.0916 | 0.0301 | -0.1507 - -0.0307 | -3.049 | **2.30e-3** |

## Discussion

An open question in the literature is whether autistic people underutilize prior experience when processing new sensory input. In this study, we investigated whether adaptation and serial choice bias, two biases induced by previous sensory input and previous perceptual decisions, respectively, are reduced in autism. To this end, we tested adolescents with and without ASD in two tasks that both used line orientation stimuli but were designed to induce either adaptation or serial choice bias. Importantly, in contrast to and in advance of previous studies, we probed adaptation and serial choice biases using the same stimuli, similar task designs, and a single sample of subjects, allowing for a more direct comparison of biases believed to arise at different stages of perceptual processing. Although we successfully induced both biases, we found no differences between the groups in the magnitude of the biases, reflecting preserved influence of previous stimuli in perception and preserved influence of previous choices in perceptual decision making, and suggesting that the past is not underutilized in autism.

Our finding that adaptation is preserved in autism may be surprising, as it conflicts with several studies that observed decreased adaptation in autism (Ewing, Leach, et al., 2013; Ewing, Pellicano, et al., 2013; Karaminis et al., 2020; Lawson et al., 2017; Pellicano et al., 2007; Turi et al., 2015; van Boxtel et al., 2016). However, there is also existing literature that has found preserved adaptation (Karaminis et al., 2015; Lawson et al., 2017; Maule et al., 2018). These diverging conclusions may be explained by the type of stimulus used in these studies. Studies that have found decreased adaptation often used complex and sometimes social stimuli, such as faces (Ewing, Leach, et al., 2013; Ewing, Pellicano, et al., 2013; Pellicano et al., 2007) or biological motion (Karaminis et al., 2020; van Boxtel et al., 2016). In contrast, studies using more simple stimuli and low-level features, such as color (Maule et al., 2018) and shapes (Karaminis et al., 2015) have found preserved adaptation. In line with this, we find preserved adaptation using line stimuli that were relatively simple and focused on a low-level feature, namely orientation. It may be that differences in adaptation arise only for complex or social stimuli, in line with the description of ASD as a social disorder. Alternatively, previously found differences in adaptation for complex and social stimuli may reflect differences in the amount of attention paid to these stimuli, as attention can boost the magnitude of adaptation (see e.g. Alais & Blake, 1999; Kreutzer et al., 2015; Lankheet & Verstraten, 1995; Raymond, 2000), and autistic individuals may attend less to social stimuli (see e.g. Simmons et al., 2009 for an overview).

With regards to the serial choice bias, we found that perceptual decisions are biased towards the previous perceptual decision, with no differences between the groups. This is not in line with expectations following the Weak Central Coherence (WCC) account, which hypothesizes that perceptual integration is impacted and from which would follow that serial choice bias may be reduced in autism (Happé & Frith, 2006), nor is it in line with the idea that an overestimation of the volatility of the environment, as found in autism (Lawson et al., 2017), may lead to a reduced leveraging of the past and thus a reduction in serial choice bias. Our finding is also in contrast with a study that has found increased influence of recent choices (Feigin et al., 2021) and with a study that found decreased influence of the past (Lieder et al., 2019).

Some previous research has investigated differences in perception as something that varies across the autism spectrum. For instance, Pellicano et al. (2007) found that adaptation magnitude was more decreased in an autistic children sample that scored higher on social atypicalities in comparison to the overall ASD group. Similarly, Lawson et al. (2017) found that adaptation magnitude decreased with autistic traits in an autistic adult sample and with autistic traits and sensory sensitivity in a non-autistic adult sample. Following this, we explored whether perceptual choice patterns varies with the severity of sensory symptomatology. Although we found some evidence of an effect, with autistic participants with more severe sensory atypicalities showing reduced influence of previous fast decisions on subsequent decisions compared to autistic participants with weaker sensory atypicalities, upon visual inspection the general linear model did not provide an adequate fit to the data, prompting caution when interpreting this finding. More research is needed to determine if there are indeed (subtle) effects.

In conclusion, we find that the use of the past is preserved in autism, suggesting that autistic individuals are able to leverage temporal context similarly to their non-autistic peers. This contradicts hypotheses that describe sensory atypicalities in autism as a result of a reduced integration of perceptual input with its temporal context.

## Acknowledgments

We extend our gratitude to our participants and their parents for supporting this research.

We also thank Christian Utzerath, Amy Abelmann, and Iris C. Schmits for assistance in recruitment (CU, AA), data collection (CU, AA, IS), and cover story development (IS).

This study was financed by the European Research Council, ERC Starting Grant 678286 CONTEXTVISION. JB has been supported by the EU-AIMS (European Autism Interventions) and AIMS-2-TRIALS programmes which receive support from Innovative Medicines Initiative Joint Undertaking Grant No. 115300 and 777394, the resources of which are composed of financial contributions from the European Union’s FP7 and Horizon2020 Programmes, and from the European Federation of Pharmaceutical Industries and Associations (EFPIA) companies’ in-kind contributions, and AUTISM SPEAKS, Autistica and SFARI; and by the Horizon2020 supported programme CANDY (Grant No. 847818).

## Supplemental materials

**Supplemental table 1:** Autism Diagnostic Interview – Revised

| ASD group member | Scales |  |  |
| --- | --- | --- | --- |
|  | Social interaction | Communication and language | Restricted and repetitive behaviors |
| 1 | 12 | 12 | 3 |
| 2 | 19 | 18 | 5 |
| 3 | 14 | 15 | 2 |
| 4 | 7 | 18 | 1 |
| 5 | 18 | 18 | 2 |
| 6 | 10 | 9 | 2 |
| 7 | 9 | 11 | 1 |
| 8 | 14 | 7 | 0 |
| 9 | 26 | 21 | 3 |
| 10 | 20 | 17 | 2 |
| 11 | 11 | 8 | 3 |
| 12 | 10 | 12 | 4 |
| 13 | 19 | 10 | 5 |
| 14 | 6 | 14 | 2 |
| 15 | 13 | 4 | 4 |
| 16 | 19 | 17 | 10 |
| 17 | 29 | 19 | 4 |
| 18 | 25 | 18 | 4 |
| 19 | 24 | 12 | 4 |
| 20 | 30 | 17 | 9 |
| 21 | 31 | 21 | 6 |
| 22 | 8 | 13 | 8 |
| 23 | 10 | 11 | 2 |
| 24 | 15 | 15 | 0 |
| 25 | 37 | 20 | 9 |
| 26 | 19 | 13 | 5 |
| 27 | 28 | 17 | 1 |
| 28 | 9 | 12 | 2 |
| 29 | 18 | 14 | 2 |
| 30 | 32 | 17 | 17 |
| 31 | 18 | 9 | 6 |

## Notes

### Competing Interest Statement

This study was financed by the European Research Council, ERC Starting Grant 678286 CONTEXTVISION. Jan K. Buitelaar has been supported by the EU-AIMS (European Autism Interventions) and AIMS-2-TRIALS programmes which receive support from Innovative Medicines Initiative Joint Undertaking Grant No. 115300 and 777394, the resources of which are composed of financial contributions from the European Union's FP7 and Horizon2020 Programmes, and from the European Federation of Pharmaceutical Industries and Associations (EFPIA) companies' in-kind contributions, and AUTISM SPEAKS, Autistica and SFARI; and by the Horizon2020 supported programme CANDY (Grant No. 847818).

